# coiaf: directly estimating complexity of infection with allele frequencies

**DOI:** 10.1101/2022.05.26.493561

**Authors:** Aris Paschalidis, Oliver J. Watson, Ozkan Aydemir, Robert Verity, Jeffrey A. Bailey

## Abstract

In malaria, individuals are often infected with different parasite strains; the complexity of infection (COI) is defined as the number of genetically distinct parasite strains in an individual. Changes in the mean COI in a population have been shown to be informative of changes in transmission intensity with a number of probabilistic likelihood and Bayesian models now developed to estimate the COI. However, rapid, direct measures based on heterozygosity or *FwS* do not properly represent the COI. In this work, we present two new methods that use easily calculated measures to directly estimate the COI from allele frequency data. Using a simulation framework, we show that our methods are computationally efficient and comparably accurate to current methods in the literature. Through a sensitivity analysis, we characterize how the bias and accuracy of our two methods are impacted by the distribution of parasite densities and the assumed sequencing depth and number of sampled loci. We further estimate the COI globally from *Plasmodium falciparum* sequencing data using our developed methods and compare the results against the literature. We show significant differences in estimated COI globally between continents and a weak relationship between malaria prevalence and COI.

**Author summary:** Computational models, used in conjunction with rapidly advancing sequencing technologies, are increasingly being used to help inform surveillance efforts and understand the epidemiological dynamics of malaria. One such important metric, the complexity of infection (COI), indirectly quantifies the level of transmission. Existing “gold-standard” COI measures rely on complex probabilistic likelihood and Bayesian models. As an alternative, we have developed the statistics and software package *coiaf*, which features two rapid, direct measures to estimate of the number of genetically distinct parasite strains in an individual (the COI). Our methods were evaluated using simulated data and subsequently compared to current “state-of-the-art” methods, yielding comparable results. Lastly, we examined the distribution of the COI in several locations across the world, identifying significant differences in COI between continents. *coiaf*, therefore, provides a new, promising framework for rapidly characterizing polyclonal infections.

## Introduction

Malaria remains a leading cause of death worldwide—in 2020, there were an estimated 241 million cases, and 627,000 deaths around the globe [1]. Despite the considerable burden of malaria, these numbers represent the substantial global progress made to control malaria in the last two decades. The WHO reports that 1.5 billion malaria cases and 7.6 million malaria deaths were averted globally from 2000 to 2019 [2]. The majority of these gains reflect an increase in vector control initiatives [3–5], the development of highly efficacious antimalarial combination therapies [6–8], and improved case management through the deployment of rapid diagnostic tests (RDTs) [9–14]. However, evidence indicates that progress has slowed and that there is a need for new approaches to capitalize on the gains already made [2].

One approach is the use of computational methods, which often rely on recent advances in genetic sequencing and provide an increased understanding of malaria biology, to help inform control efforts [15–17]. For example, molecular and genomic epidemiology surveillance tools, which have been developing rapidly, can be used to track drug resistance and understand and evaluate control efforts [18, 19]. Moreover, computational methods have been applied in the identification of polyclonal infections—malaria infections with multiple distinct strains [20, 21]. Such polyclonal infections introduce additional genetic complexity that is often difficult to account for computationally. As a result, many of the population genetic tools often applied for the study of other organisms are unsuitable for studying malaria, and researchers often rely on limiting genetic analyses to individuals who are monoclonally infected [22].

Although determining the most informative metrics is an active field of investigation [23], one important metric is the complexity of infection (COI). Sometimes referred to multiplicity of infection, although this is generally reserved for infections within the same cell, the COI represents the number of genetically distinct malaria genomes or strains that can be identified in a particular individual. These polyclonal infections may arise from: (*i*) a single infectious mosquito feeds on a human host, transferring several genetically distinct parasite strains (often referred to as a co-transmission event [24, 25]) or (*ii*) two or more infectious mosquitoes with distinct malaria strains feed on an individual (known as a superinfection event [25, 26]). Measures of genetic diversity and the COI are increasingly used for inferring malaria transmission intensity and evaluating malaria control interventions [19]. Transmission intensity has been shown to impact the contribution of each event towards the generation of within-host parasite genetic diversity [23]. Superinfection is modulated by the host’s current infections [27], age, and exposure acquired immunity [28]. Additionally, the COI provides a practical approach for identifying monoclonal infections to simplify genomic analyses.

Traditionally, the COI was measured in one or a few regions of the genome, relying on the enumeration of the maximal number of haplotypes detected through PCR amplification at genes or markers encoding highly diverse length polymorphisms. Two of the most common markers are the merozoite surface proteins 1 and 2 (*msp1* and *msp2*), surface proteins found on the merozoite stage of the malarial parasite [29, 30]. Traditional methods are hindered by limitations on the number of loci examined [31, 32], and lack the ability to detect parasites at low parasitaemia in samples [33]. Furthermore, sanger sequencing lacks the sensitivity for low parasitemia samples without employing laborious subcloning [34].

High-throughput sequencing provides more sensitive and specific methods. As a result, new computational methods have been developed to determine the COI. Two early proposed methods were the *FwS* metric by Aubern et al., which characterizes within-host diversity and its relationship to population-level diversity [31], and the *estMOI* software, which utilizes local phasing information of microhaplotypes within read pairs to estimate the COI [33]. Unfortunately, the *FwS* metric does not directly relate to the COI nor have a concrete biological interpretation, and the *estMOI* software relies on observed local haplotypes and heuristic interpretation. More recently, new tools been developed to better measure the COI beyond the maximal observed haplotype. For instance, the *DEploid* software package uses haplotype structure to infer the number of strains, their relative proportions, and the haplotypes present in a sample [35]. However, it is known that *DEploid* under predicts COI for high COI infections [25]. Other methods have been developed to examine the relatedness between parasite strains [36, 37]. The current “state-of-the-art” method for determining the COI of a sample is *THE REAL McCOIL*, which is an extension of *COIL* method [32]. *THE REAL McCOIL* employs a Bayesian approach, turning heterozygous data into estimates of allele frequency using Markov chain Monte Carlo methods and jointly estimating the likelihood of the COI [38].

Despite various methods for estimating the COI, no rapid, direct measures have been developed to work effectively on a set of loci or at the genome-wide level. In this work, we present two new methods that use easily calculable metrics to directly estimate the COI from allele frequency data. Our two methods closely resemble the categorical and proportional methods implemented in *THE REAL McCOIL* [38], yet are geared towards large numbers of loci and providing rapid estimates of COI. Using our methods, we have developed the software package *coiaf* in the programming language 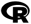, a language and environment for statistical computing and graphics [39].

## Materials and methods

### Problem formulation

Current “state-of-the-art” approaches for estimating the COI rely on identifying the number of different parasites present in an infection using high-throughput sequencing. In monoclonal infections, i.e., infections composed of only one parasite strain (COI = 1), all sequence reads should be identical, originating from the same parasite strain. However, in mixed infections, a combination of each parasite strain, proportional to the strain’s abundance, will contribute to the observed sequence reads. At genetic loci containing variation, there is an increased chance of observing multiple alleles as the number of unrelated parasite strains within an infection increases. Therefore, the likelihood of observing multiple alleles at any locus is dependent on the number of parasite strains in an infection and the prevalence of genetic polymorphisms in the population.

We focus on only biallelic SNPs—the vast majority of loci—and define the major allele as the allele that is most prevalent in a population. We note that any multiallelic site can be collapsed to a biallelic site, although information will be lost. Assuming for any individual there are *l* biallelic loci, we define the within-sample allele frequency (WSAF) as a vector **ŵ** of length *l* composed of the frequencies of the reference allele at each locus for a single individual infection and the population-level allele frequency (PLAF) as a vector 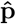 of length *l* composed of the frequencies of the reference allele at each locus across a population. The PLAF may be represented as the mean of the WSAF across a population, i.e., 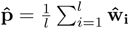.

We further define the population-level minor allele frequency (PLMAF) as a vector **p** of length *l* composed of the frequencies of the minor allele at each locus across a population, namely **p** = (*p*_1_, …, *p*_*l*_), where *p*_*i*_ ∈ [0, 0.5]. The within-sample minor allele frequency (WSMAF) is, additionally, defined as a vector **w** of length *l* composed of the frequencies of the population-level minor allele at each locus for a single individual infection. For instance, the WSMAF will be equal to one when all sequence reads observed at a given locus are of the population-level minor allele. Given the WSAF and the PLAF, the WSMAF and the PLMAF may be expressed as follows:

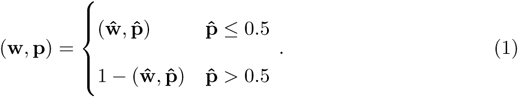

### Variant and Frequency Methods

Our overall goal is to estimate the COI of the sample, denoted by *k*, using the WSMAF and the PLMAF. We do this by comparing observed data to derived expressions that define a relationship between the WSMAF, the PLMAF, and the COI. We present two alternative expressions that we refer to as the Variant Method and the Frequency Method.

In the Variant Method, we examine a set of SNPs and express the probability of SNP *i* being heterozygous with respect to the PLMAF and the COI. We define *V*_*i*_, a Bernoulli random variable that takes the value of one if a site is heterozygous and zero otherwise. The probability that locus *i* is heterozygous, written as ℙ (*V*_*i*_ = 1), will be equal to one minus the probability that a locus is homozygous (see Appendix A in S1 Appendices). We thus write,

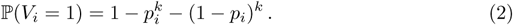

As the COI increases, the probability of observing a heterozygous locus within an infection also increases (see Fig 1). We note that this is the same expression used within the Categorical method of *THE REAL McCOIL* (Eq (2)) [38].

**Fig 1.**
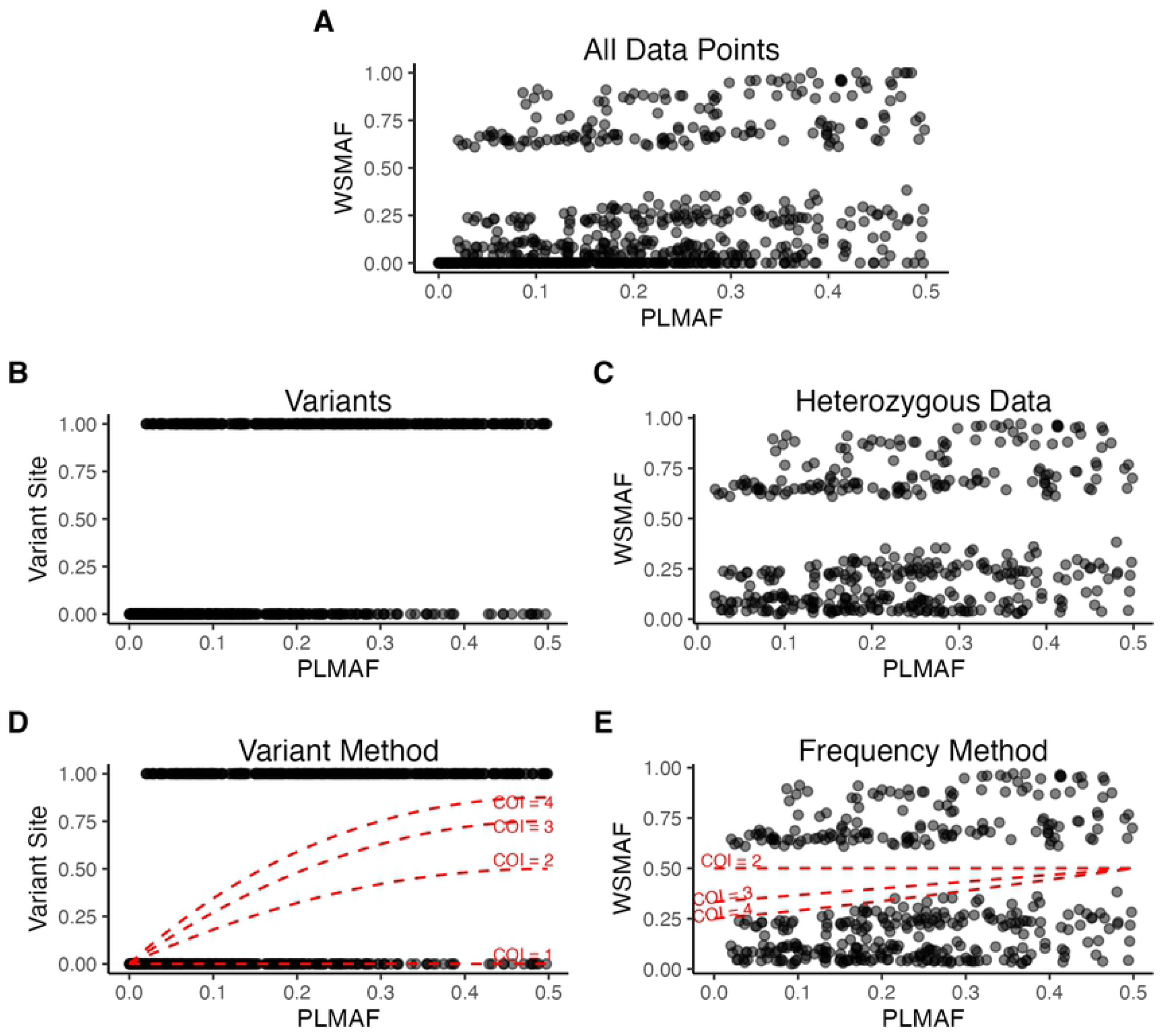
Flowchart of methods. In **A**, the relationship between the WSMAF and the PLMAF is shown for an example simulation with a COI of 4. In **B**, data have been processed so that loci are deemed variant if they are heterozygous and invariant otherwise. In **C**, homozygous data have been filtered out. Following processing of data, Eq (2) and Eq (3) have been plotted for varying COIs from 1 to 4 in **D** and **E**, respectively.

In the Frequency Method, we focus on the expected value of the within-sample minor allele frequency. For the sake of simplicity, the complete derivation has been left to Appendix A in S1 Appendices. Briefly, we determine the probability of a particular strain carrying the minor allele and then determine the expected WSMAF by summing over the expected contribution of each strain. We represent the expected value of the within-sample minor allele frequency given that a site is heterozygous as follows:

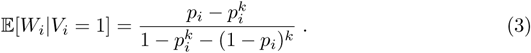

### Estimation method

Given data, *D* : {(*p*_*i*_, *w*_*i*_, *d*_*i*_), *i* = 1, …, *l*}, where *p*_*i*_ is the PLMAF at locus *i, w*_*i*_ is the WSMAF at locus *i*, and *d*_*i*_ is the coverage at locus *i*, we next explore our methods to approximate the COI of a sample. Data are first processed to account for sequencing error. This process denotes loci at which there was suspected sequencing error as homozygous instead of heterozyogus (for additional information, see Appendix C in S1 Appendices).

Following adjustment for sequence error, we consider an arbitrary data point (*p*_*i*_, *w*_*i*_, *d*_*i*_). Recall that the Variant Method and the Frequency Method examine different random variables. Specifically, the Variant Method identifies the probability of a locus being heterozygous, ℙ(*V*_*i*_ = 1), and the Frequency Method identifies the expected value of the WSMAF given a site is heterozygous, 𝔼[*W*_*i*_|*V*_*i*_ = 1]. In order to determine the COI, we utilize Eq (2) and Eq (3). We solve the following minimization problem for the Variant Method:

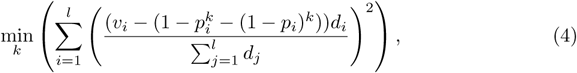

where *v*_*i*_ is defined as the value of the random variable *V*_*i*_ and *q* ≥ 1. Note that 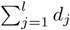 is a constant and, therefore, we may remove it from our expression to get our final minimization problem:

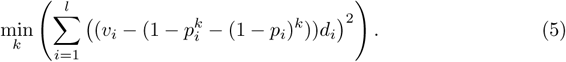

Similarly, we solve the following minimization problem for the Frequency Method:

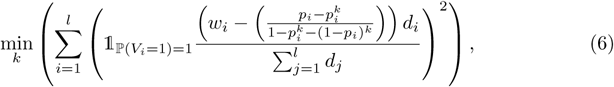

where *q* ≥ 1. Once again, we may remove the constant from our expression to get our final minimization problem:

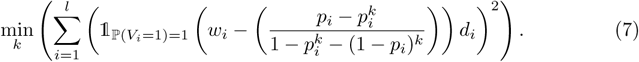

Note that the estimation methods described minimize the sum of squared residuals between the observed data and the relationships derived in Eq (2) and Eq (3).

### Solution methods

We solve this optimization problem using two methods: (*i*) assuming discrete values of the COI and (*ii*) assuming continuous values of the COI. Recall that the COI is defined as the number of genetically distinct malaria parasite strains an individual is infected with. While a continuous value of the COI has no direct biological interpretation, significant departures from discrete values are expected due to parasite relatedness, and a range of other biological phenomenon including, but not limited to, overdispersion in sequencing and parasite densities. Therefore, a continuous value of the COI may provide a more accurate representation of the overall population of samples being studied. Furthermore, as relatedness in mixed infections is common [37], a continuous COI may be able to provide insights into the degree of relatedness between parasite strains in mixed infections and detect highly-related polyclonal infections that may traditionally be categorized as monoclonal.

To solve the discrete versions of the previously defined optimization problems we use a brute force approach, which involves computing the objective function for each COI considered. As brute force approaches can be computationally inefficient, we limit the range of values of the COI. To solve the continuous versions of the optimization problems, we utilize 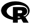 built-in optimization function [39]. In particular, we leverage a quasi-Newton L-BFGS-B approach with box constraints [40]. We set the lower and upper bounds of the COI as 1 and 25, respectively, with the default starting value of the COI equal to two. Note that in both the discrete and continuous case, the upper bound of the COI is much larger than most COIs seen in the real world [38, 41]. In both cases, we also provide the capability to determine the 95% confidence interval for our COI estimates by leverage bootstrapping techniques [42] (see Appendix E in S1 Appendices for more details).

### Data

To evaluate the accuracy and sensitivity of our methods, we created a simulator that generates synthetic sequencing data for a number of individuals in a given population. In overview, each individual is assigned a COI value. The haplotype of each strain is then assigned by sampling from the population level minor allele frequency. Next, we simulate the number of sequence reads mapped to the reference and alternative allele by sampling in proportion to the parasite densities for each strain. After simulating sequence error, the mapped sequence reads are then used to derive the within-sample minor allele frequency. A detailed description of our simulator can be found in Appendix F in S1 Appendices.

In addition to simulated data, we use sequencing data sampled from infected individuals worldwide to compare our methods to the current state-of-the-art COI estimation metric and to investigate the distribution of the COI across the world. We analysed over 7,000 *P. falciparum* samples from 28 malaria-endemic countries in Africa, Asia, South America, and Oceania from 2002 to 2015 from the MalariaGEN *Plasmodium falciparum* Community Project [43]. Detailed information about the data release including brief descriptions of contributing partner studies and study locations is available in the supplementary of MalariaGEN et al. [43]. We used the provided variant call files (VCFs) generated using a standard analysis pipeline. The median read depth of coverage of the initially sequenced field isolates was 73 across across all samples. After removing replicate samples, mixed-species samples, and samples with a low coverage, suspected contamination or mislabelling, 5,970 samples remained for further analysis. Genomic data were further filtered for high quality biallelic coding and non-coding SNPs as outlined in Zhu et al. [37]. Additionally, data were filtered to sites that are part of the core genome.

To apply our developed methods, we must estimate the population-level frequency of the minor allele. Consequently, we sought to assign the samples to a suitable number of geographic regions such that the number of samples per region was suitable for the reliable estimation of the population-level minor allele frequencies. We used the Partitioning Around Medoids (PAM) algorithm to solve a k-medoids clustering problem [44, 45] to group samples based on the longitude and latitude of sample collection. We next calculated the silhouette information for each clustering of k groups [46], arriving at 24 regions globally (see Appendix K.2 in S1 Appendices for a map of locations). Given these 24 clusters, we filtered SNPs to variants with a population level alternative allele frequency greater than 0.005 in each region. The 0.005 frequency cutoff was chosen as sequence error likely obscures the detection of true variation from parasite strains comprising less than 0.5% of total parasite density. Clusters of data were additionally traced to a specific continent and subregion as defined by the World Development Indicators [47, 48].

## Results

### Performance on simulated data

Using our simulator, we simulated data for 1,000 loci with a read depth of 100 at each locus. Data was simulated with a complexity of infection ranging from 1 to 20. This simulation did not introduce error in order to determine optimal performance based on sampling. Our methods, therefore, accounted for no sequencing error. The results of running the discrete version of both the Variant Method and the Frequency Method are illustrated in Fig 2A and Fig 2B, respectively.

**Fig 2.**
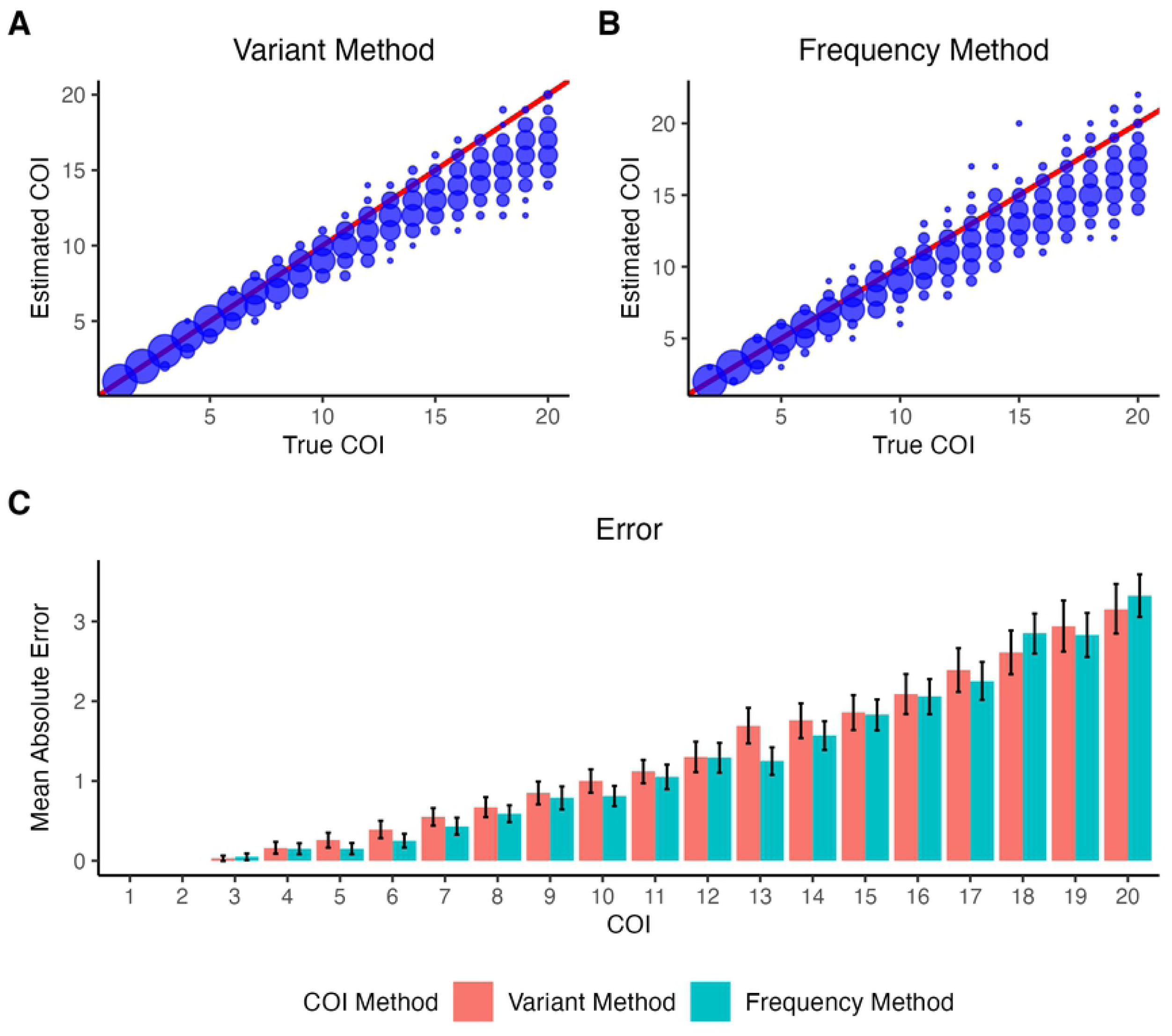
Estimating the COI on simulated data. The performance of the Variant Method (**A**) and Frequency Method (**B**) is shown for 100 simulations of a COI of 1-20 with 1,000 loci, a read depth of 100, no error added to the simulations, and no sequencing error assumed. Point size indicates density, with the red line representing the line *y* = *x*. The mean absolute error for each method is shown in **C**. The black bars indicate the 95% confidence interval.

The Variant Method and the Frequency Method both perform well for all COIs between 1 and 20. Notably, the lower the true value of the COI, the better our models perform with a mean absolute error close to zero. As the COI increases, our models exhibit more variability across subsequent iterations and underestimate the true COI. For example, at a COI of 20, the estimated COI from the Frequency Method ranges from 14 to 22—the maximum COI our model could output was 25 in these trials. Nevertheless, the majority of predicted COIs remain close to the true COI as witnessed by the low mean absolute errors of at most 3.32 in Fig 2C. Comparing our two methods, we find that the Variant Method and the Frequency Method perform equally with insignificantly different mean absolute errors (p-value = 0.396) and biases (p-value = 0.661).

### Sensitivity analysis

In order to understand the sensitivity of our models to alterations in the parameters considered, we tested the performance of the discrete and continuous representations of the Variant Method and the Frequency Method, assessing changes in the accuracy of our predictions. For each sample, we utilized bootstrapping techniques [49, 50] to determine the mean absolute error and bias of the predicted COI compared to the true COI. Furthermore, we ran each algorithm several times to ensure reliable results. A description of several key parameters perturbed and their default values can be found in Appendix G in S1 Appendices. Resulting figures can be found in Appendix K.1 in S1 Appendices.

Here, we highlight the effect of varying two metrics that can be controlled in the field: the read depth at each locus and the number of loci sequenced. Sequencing more loci at larger read depths is preferred as this results in more higher quality data. In general, as the coverage at each locus increases, the performance of our methods also improves (see Appendix K.1 Fig 2 and Fig 3 in S1 Appendices). This relationship, however, is not linear. Rather, a non-monotonic relationship is observed between sequence coverage and the mean absolute error, with diminishing returns in performance observed with sequence coverage greater than 100. As was the case for our coverage data, when the number of loci sequenced is low, around 100 loci, our methods have a high variability and tend to underpredict the true COI (see Appendix K.1 Fig 4 and Fig 5 in S1 Appendices). However, as the number of loci increases to 1,000, the performance increases. In addition, increasing the number of loci examined above a certain threshold, in this case 1,000 loci, does not seem to substantially impact the performance of our models. However, note that an increase in the number of loci does reduce the variance of our estimates.

**Fig 3.**
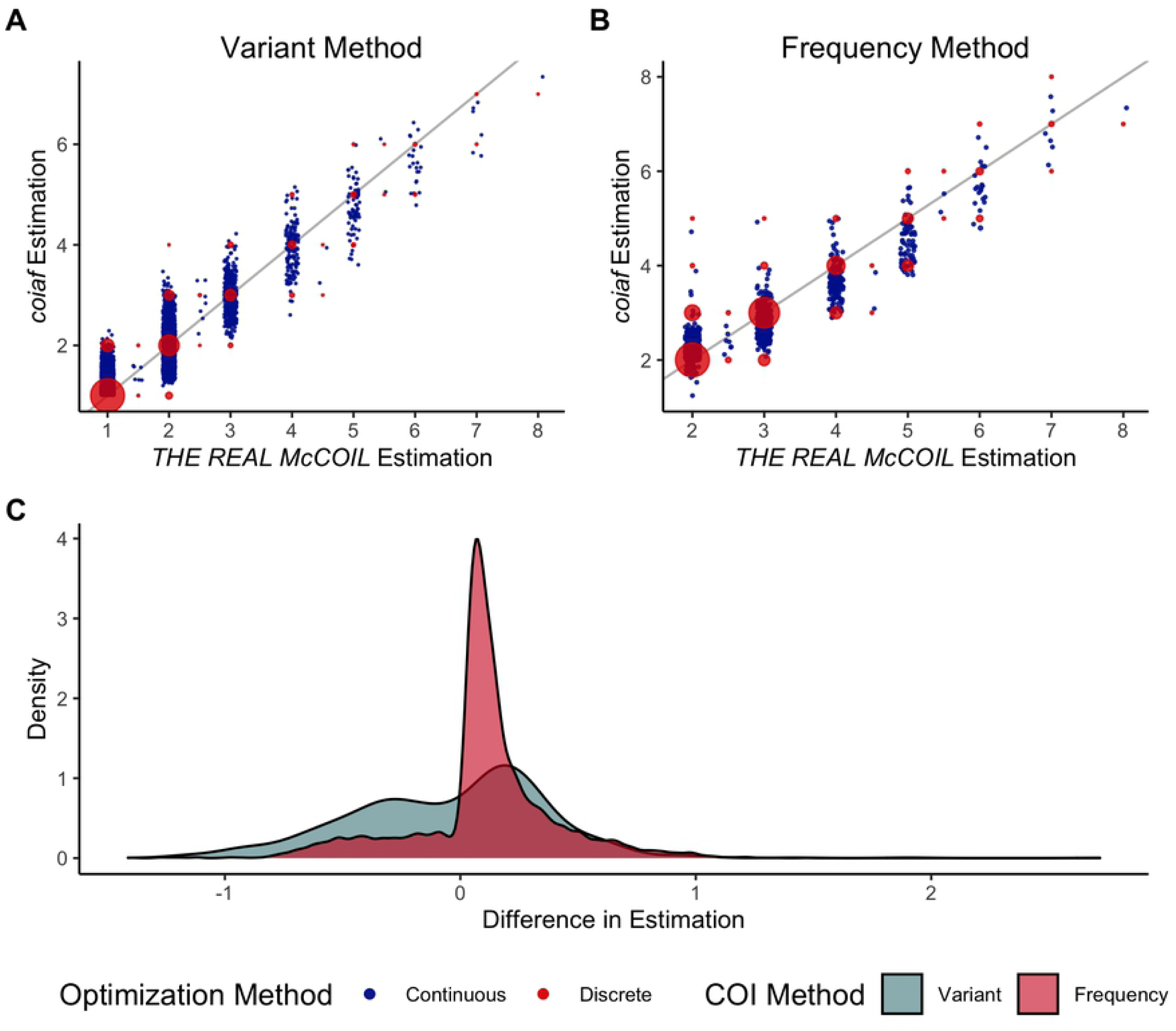
Comparison Between *THE REAL McCOIL* and *coiaf*. The COI estimation using (**A**) the Variant Method and (**B**) the Frequency Method is compared against the *THE REAL McCOIL*. In **C** the distribution of differences between our estimation and the *THE REAL McCOIL*’s estimation is shown.

**Fig 4.**
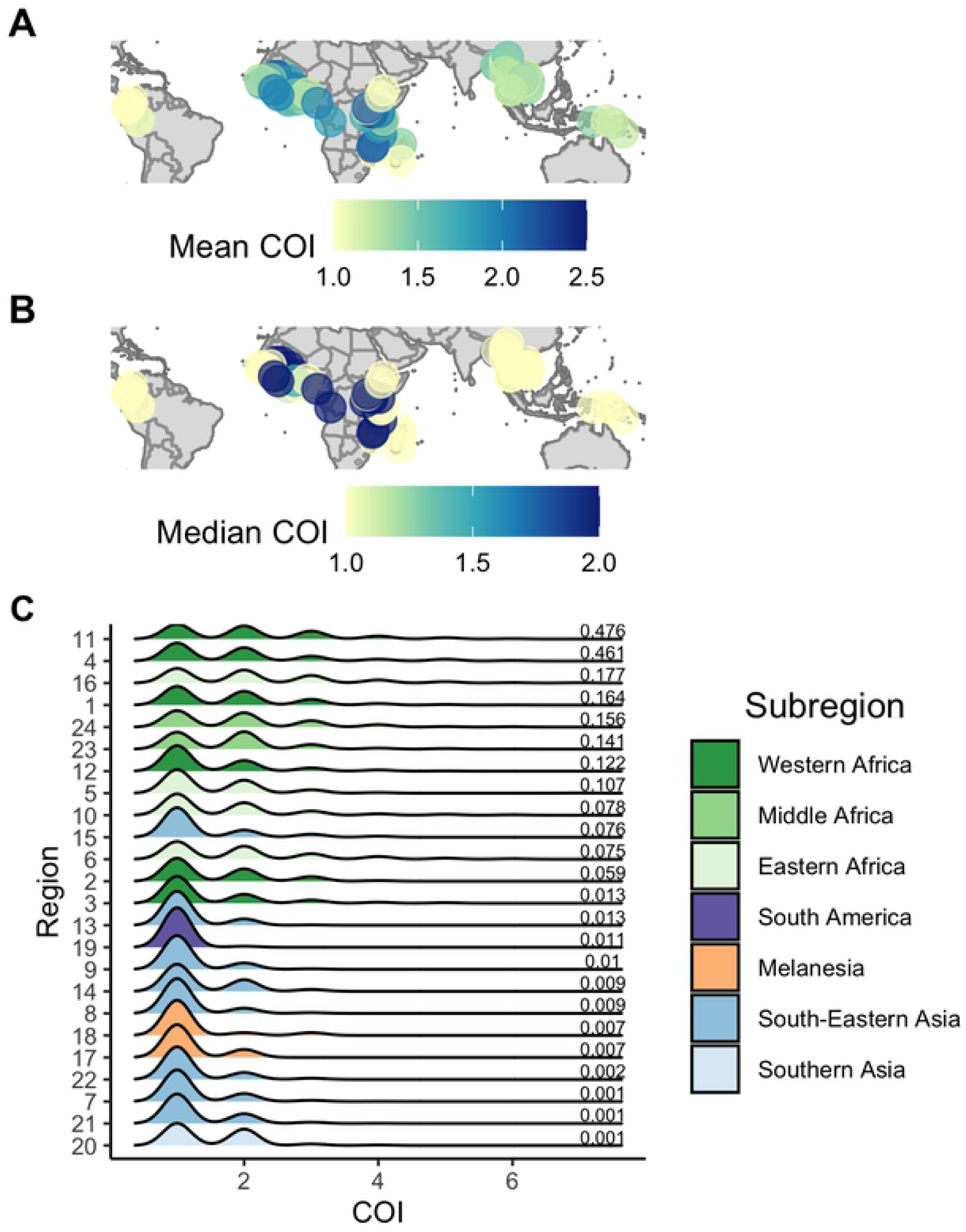
COI Across the Globe. In **A** we plot the mean COI of all samples in each study location within the 24 regions, in **B** we plot the mean COI. The color and size of each point represents the magnitude of the COI. In **C**, we draw a density plot for each region, where the color of the plot indicates in what subregion the data was sampled. The plots are sorted by the median microscopy prevalence in children aged 2-10 years old as estimated in the Malaria Atlas Project [9, 10, 52] and indicated to the right of each density plot.

### Comparison to “state-of-the-art” methods

In this section we compare our novel methods to the current “state-of-the-art” method used to estimate the COI, *THE REAL McCOIL* [38]. As was previously described, we grouped our data into 24 regions around the world. To estimate the COI for each of the 5,970 samples, we examined an average of 32, 362 (range: 15, 276-40, 272) loci in each region (see Appendix I in S1 Appendices). Furthermore, we ran five repetitions of the *THE REAL McCOIL* on each sample, with a burn-in period of 1, 000 iterations followed by 5, 000 sampling iterations, and using standard methodology to confirm convergence between Monte Carlo Markov chains [51]. For additional information, see Appendix J in S1 Appendices. Fig 3 examines the COI estimation of *THE REAL McCOIL* and *coiaf* for all samples. We note that the relationship between *coiaf* ‘s estimated COI and *THE REAL McCOIL*’s estimated COI for each of the 24 individual regions is shown in Appendix K.3 Fig 1 and Fig 2 in S1 Appendices.

We observe that the Variant Method and the Frequency Method are strongly correlated with the estimates from *THE REAL McCOIL*. In (Fig 3A and Fig 3B), When the COI is estimated to be below five, both methods estimate COI values that are close to one another. However, as the estimated COI increases, there is a greater variability in predictions. At these high COI values, our methods tend to estimate the COI within three of the true COI (Fig 3C). As expected, the continuous estimation methods align with the discrete estimation methods. Furthermore, we note that the Frequency Method does not show estimates for when *THE REAL McCOIL* predicts a COI of one. This is because the Frequency Method at a COI of one is undefined; at a COI of one, there would be no heterozygous loci, which are used in the Frequency Method. To quantify the relationship between our novel software and the current “state-of-the-art” method, we fit linear regression models to the data and evaluated the Pearson correlation between estimation methods. The results are reported in Table 1 and indicate that each of the methods introduced in *coiaf* is highly correlated with *THE REAL McCOIL*.

**Table 1.**
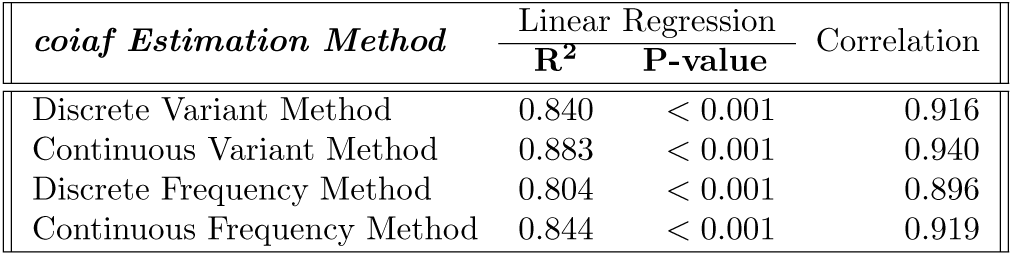
Relationship Between *coiaf* and *THE REAL McCOIL*. A linear regression model was fit to the data to evaluate the relationship between *coiaf* ‘s and *THE REAL McCOIL*’s estimation method. Furthermore, the Pearson correlation between the estimated COI’s was computed.

| *coiaf* Estimation Method | Linear Regression |  | Correlation |
| --- | --- | --- | --- |
|  | R<sup>2</sup> | P-value |  |
| Discrete Variant Method | 0.840 | < 0.001 | 0.916 |
| Continuous Variant Method | 0.883 | < 0.001 | 0.940 |
| Discrete Frequency Method | 0.804 | < 0.001 | 0.896 |
| Continuous Frequency Method | 0.844 | < 0.001 | 0.919 |

### Mapping COI worldwide

To demonstrate *coiaf*’s utility and better understand global patterns of the COI, we examined the distribution of COI in twenty-four different regions. In each region we studied an average of 248 (range: 29 to 909) samples and 32,362 loci (range: 15,276 to 40,272) (see Appendix I in S1 Appendices). Our samples can be traced to four different continents, with most originating in either Africa (55.5%) or Asia (41.8%). All other samples were sequenced in Oceania (2.03%) or the Americas (0.620%).

Fig 4 highlights the mean and median COI across all samples in each of the 24 regions outlined previously. We, furthermore, aimed to understand the relationship between the complexity of infection and malaria prevalence by leveraging estimates of malaria microscopy prevalence in children aged 2-10 years old generated by the Malaria Atlas Project [9, 10, 52]. Fig 4C plots the density of the COI for each region sorted by the region’s malaria prevalence. Table 2 outlines the mean COI in each of the four continents and seven subregions analyzed.

**Table 2.** Mean COI. Mean COI across each continent and subregion analyzed.

| Continent | Subregion | Number of Samples | Mean COI (SD) |
| --- | --- | --- | --- |
| Africa | Eastern Africa | 739 | 1.88 (1.10) |
| Africa | Middle Africa | 579 | 1.87 (0.948) |
| Africa | Western Africa | 1,996 | 1.87 (1.05) |
| Americas | South America | 37 | 1.03 (0.164) |
| Asia | South-Eastern Asia | 2,421 | 1.25 (0.506) |
| Asia | Southern Asia | 77 | 1.52 (0.641) |
| Oceania | Melanesia | 121 | 1.20 (0.440) |

Of all the continents, Africa has the highest mean COI of 1.87, followed by Asia with a mean COI of 1.26, Oceania with a mean COI of 1.20, and the Americas with a mean COI of 1.03. A Nemenyi post-hoc test [53] indicates that while Africa is statistically different than all the other continents (p-value: *<* 0.001 in all cases), there exist no significant differences between each pairing of the other three continents (Americas vs Asia p-value: 0.204, Americas vs Oceania: p-value: 0.568, p-value: Asia vs Oceania: 0.816). Within each continent, there exist further differences among the subregions. No statistically significant difference was found between each of the three subregions in Africa (Eastern vs Middle p-value: 0.914, Eastern vs Western p-value: *>* 0.999, Middle vs Western p-value: 0.886). However, in Asia, there was a statistically significant difference between the mean COI in South-Eastern Asia and Southern Asia (p-value: 0.0282).

We found a positive correlation between the COI and the microscopy prevalence (Table 3). In regions with a lower prevalence, there were few samples with a COI larger than two. In fact, in regions with a prevalence less than or equal to 0.01, more than 95% of samples had a COI of one or two. In regions with a higher prevalence, there were more samples with a higher COI—in regions where the prevalence was greater than or equal to 0.1, more than 20% of samples had a COI greater than two.

**Table 3.** Relationship Between *coiaf* and malaria prevalence. A linear regression model was fit to the data to evaluate the relationship between the COI and prevalence. Furthermore, the Pearson correlation was examined.

| *coiaf* Estimation Method | Linear Regression |  | Correlation |
| --- | --- | --- | --- |
|  | R<sup>2</sup> | P-value |  |
| Discrete Variant Method | 0.0770 | $< 0.001$ | 0.278 |
| Continuous Variant Method | 0.0787 | $< 0.001$ | 0.281 |
| Discrete Frequency Method | 0.0190 | $< 0.001$ | 0.138 |
| Continuous Frequency Method | 0.0218 | $< 0.001$ | 0.148 |

## Discussion

Despite advances in sequencing technologies and the development of various methods for estimating the COI, no direct measures have been developed to rapidly estimate the COI. Here we develop two direct measures based on minor allele frequencies that address these needs. Our methods can provide rapid estimates of the COI in under a second. When compared to the current state-of-the-art COI estimation software, *THE REAL McCOIL, coiaf* was able to estimate the COI with similar accuracy approximately 250 times faster (see Appendix H in S1 Appendices) and required more than eight times less memory. Moreover, our methods can be scaled to fully use all available SNPs within the genome and provide a continuous measure that can provide insight into relatedness.

Through a number of simulations, we further explored how changing key sequencing variables, such as the number of loci and read depth at each locus, altered our software’s performance. We showed that for samples with a low and moderate COI our methods were able to accurately predict the COI even with a low coverage and number of loci, however, as the COI increased, these parameters became more important—a lack of sufficient sequencing corresponded with underprediction of the true COI. Additionally, we demonstrated that there are several important factors that can greatly influence results, such as sequencing error or overdispersion in parasite density. Importantly, we also show that the population mean WSMAF is an unbiased estimator of the PLMAF (see Appendix B in S1 Appendices). This provides further advantages to using allelic read depth for COI estimation rather than haplotype calls, which are known to lead to biased estimates of population allele frequency if the COI of samples is not accounted for [54].

An application of *coiaf* on several thousand *P. falciparum* samples from malaria-endemic countries in four continents from 2002-2015 [43] resulted in a comprehensive map of the complexity of infection worldwide. This study builds on previous reviews of the distribution of the COI globally [55], and is the first study to our knowledge to provide a holistic view of the COI based on allelic read depth as opposed to traditional methods leveraging *msp1* and *msp2* haplotyping.

In general, our results were in agreement with previously reported findings. For instance, in areas with historically lower malaria prevalence, such as South America and Southern Asia, we estimated lower average COI values. In particular, in the Americas and Asia we report a mean COI of 1.03 and 1.26, respectively. Previous efforts to estimate the COI in these region have also found similarly low values of the COI. While directly comparing our estimates against these would be incorrect as the date and location of sample collection are very different, we are encouraged by the similarity in estimates of the COI. For example, in Brazil, the COI has been previously measured as low as 1.1 in the early 1990s [56]. In Papua New Guinea [41, 57, 58], Bangladesh [59] and Malaysia [60] previous estimates of COI range between 1.00 and 2.12, 1.22 and 1.58 and 1.20 and 1.37 respectively. In Africa, on the other hand, we were surprised to find little to no difference in average COI estimates between the three subregions we studied: Eastern Africa, Middle Africa, and Western Africa, despite large differences in average malaria prevalence in these regions. In contrast, previous studies of the COI in African settings have found both higher average COI values in some regions and, in general, greater variability. For example, in Cameroon large mean COIs between 2.33 to 3.82 have been reported [61–63]. Conversely, in Ghana the mean COI has been reported to be between 1.13 and 1.91 in 2012-2013 [64].

Much of the work surrounding the complexity of infection is motivated by the fact that the COI has been proposed as an indicator of transmission intensity. Unfortunately, the relationship between the COI and malaria prevalence remains an area of much debate, with many individual studies finding different relationships between the COI and malaria prevalence [65, 66]. Lopez and Koepfli report in their review article that across the 153 studies examined there was a weak correlation between mean COI and prevalence, an observation that agrees with our findings [55]. As previously noted, there are multiple patient-level factors (e.g., age and clinical status) which may affect the relationship between malaria prevalence and the COI. Additionally, Karl et al. suggested that this weak correlation may be attributed to spatial effects and the existence of geographic “hotspots” where transmission may be much higher than in surrounding areas, causing individuals to have a greater COI [67]. Moreover, multiple studies have highlighted that seasonality affects observed COI [68, 69]. Consequently, while there is undoubtedly a relationship between transmission intensity and COI, it is important to be aware of how many factors (age, clinical status, seasonality, spatial effects, parasite density, time since last infection, methods of detecting multiple infections) may impact this relationship. For example, while analysing the data from the MalariaGEN *Plasmodium falciparum* Community Project [43], metadata is only available regarding the year and location of sample collection. Without being able to account for the other factors known to impact COI, we are unable to make more meaningful interpretations of our analysis of COI patterns. Lastly, it is worth recalling that malaria prevalence is not directly related to transmission intensity. For example, two regions with the same malaria prevalence will likely have different transmission intensities if intervention coverage differs between the region. Therefore, the variance observed between malaria prevalence and the COI may reflect that malaria prevalence is itself an imperfect predictor of transmission intensity.

Our work is not without limitations. In part, limitations stem from the fact that our methods rely on certain biological assumptions, which may not be met in the real world. Additionally, the accuracy of our algorithms is impacted by sequence error. While this was not an issue in our analysis of the MalariaGEN *Plasmodium falciparum* Community Project (see Appendix K.3 Fig 5 in S1 Appendices), high levels of sequence error need to be monitored and accounted for. While our software package does allow users to account for this by providing a level of suspected sequencing error, sequence error is unlikely to be constant across the genome and accurate inference of sequencing error is itself an active research challenge [70]. Furthermore, while our methods do account for the coverage at each locus, if there is a low overall coverage for a sample, our results may underpredict the true COI. Lastly, our methods assume that the population level minor allele frequency (PLMAF) is well captured by the samples provided. Sampling bias or undersampling in a region may result in an inaccurate PLMAF estimation, which may influence our estimated COI.

In conclusion, we developed two direct measures for estimating the COI given the within sample allele frequency of a sample and the population level allele frequency. Our methods were able to estimate the COI for samples in less than a second and were shown to be accurate when compared to simulated data and current COI estimation techniques. Our software will aid in the estimation of the complexity of infection, an increasingly important population genetic metric for inferring malaria transmission intensity and evaluating malaria control interventions [19, 71].

## Supporting information

**S1 Appendices. Appendices**. Includes the complete problem formulation including derivations of the Variant Method and the Frequency Method, additional information about our methods, and all omitted figures.

## Acknowledgments

We thank the MalariaGEN *Plasmodium falciparum* Community Project for maintaining a large collection of sequencing data and variant calls.

